# Snake venom phospholipases A2 possess a strong virucidal activity against SARS-CoV-2 in vitro and block the cell fusion mediated by spike glycoprotein interaction with the ACE2 receptor

**DOI:** 10.1101/2021.01.12.426042

**Authors:** Andrei E. Siniavin, Maria A. Nikiforova, Svetlana D. Grinkina, Vladimir A. Gushchin, Vladislav G. Starkov, Alexey V. Osipov, Victor I. Tsetlin, Yuri N. Utkin

**Affiliations:** Department of Molecular Neuroimmune Signalling, Shemyakin-Ovchinnikov Institute of Bioorganic Chemistry, Russian Academy of Sciences, Moscow, 117997, Russia; N.F. Gamaleya National Research Center for Epidemiology and Microbiology, Ivanovsky Institute of Virology, Ministry of Health of the Russian Federation, Moscow, 123098, Russia

**Keywords:** antiviral activity, SARS-CoV-2, COVID-19 pandemic, phospholipase A2, coronavirus, snake venom

## Abstract

A new coronavirus was recently discovered and named severe acute respiratory syndrome coronavirus 2 (SARS-CoV-2). In the absence of specific therapeutic and prophylactic agents, the virus has infected almost hundred million people, of whom nearly two million have died from the viral disease COVID-19. The ongoing COVID-19 pandemic is a global threat requiring new therapeutic strategies. Among them, antiviral studies based on natural molecules are a promising approach. The superfamily of phospholipases A2 (PLA2s) consists of a large number of members that catalyze the hydrolysis of phospholipids at a specific position. Here we show that secreted PLA2s from the venom of various snakes protect to varying degrees the Vero E6 cells widely used for the replication of viruses with evident cytopathic action, from SARS-CoV-2 infection PLA2s showed low cytotoxicity to Vero E6 cells and the high antiviral activity against SARS-CoV-2 with IC_50_ values ranged from 0.06 to 7.71 μg/ml. Dimeric PLA2 HDP-2 from the viper *Vipera nikolskii*, as well as its catalytic and inhibitory subunits, had potent virucidal (neutralizing) activity against SARS-CoV-2. Inactivation of the enzymatic activity of the catalytic subunit of dimeric PLA2 led to a significant decrease in antiviral activity. In addition, dimeric PLA2 inhibited cell-cell fusion mediated by SARS-CoV-2 spike glycoprotein. These results suggest that snake PLA2s, in particular dimeric ones, are promising candidates for the development of antiviral drugs that target lipid bilayers of the viral envelope and may be good tools to study the interaction of viruses with host cell membranes.

## Introduction

In December 2019, a rapidly spreading community-acquired pneumonia was discovered in Wuhan (Hubei Province, China) and subsequent studies have shown that this disease is caused by a virus belonging to the coronavirus family, this conclusion soon confirmed by sequencing the full-length genome from samples (bronchoalveolar lavage fluid) of patients with pneumonia ^1^. In February 2020, WHO (https://www.who.int/dg/speeches/detail/who-director-general-s-remarks-at-the-media-briefing-on-2019-ncov-on-11-february-2020) has officially named this disease «COVID-19» (coronavirus disease 2019) and the causative virus was designated as SARS-CoV-2 (severe acute respiratory coronavirus 2) by the International Committee on Taxonomy of Viruses ^2^. As of March 2020, COVID-19 has spread worldwide and WHO has declared a global pandemic ^3^. At present the total number of infected has reached about 100 million of which nearly 2 million have died. The information is available about several vaccines which passed the third stage ^4^, but there are no drugs with experimentally proven efficacy in the treatment of coronavirus infection, which shows an urgent need for search of drugs against COVID-19.

Coronaviruses are enveloped RNA viruses form the *Coronaviridae* family which includes four genera: α-, β-, γ- and δ-coronaviruses ^5^. Of the seven known coronaviruses that infect humans, HCoV-229E and HCoV-NL63 belong to α-coronaviruses, while HCoV-HKU1, HCoV-OC43, MERS-CoV, SARS-CoV, and SARS-CoV-2 are β-coronaviruses ^6^. Coronaviruses HCoV-229E, HCoV-NL63, HCoV-HKU1 and HCoV-OC43 cause mild disease symptoms similar to the common cold ^7^. However, MERS-CoV, SARS-CoV and SARS-CoV-2 cause more severe disease associated with pneumonia, acute respiratory distress syndrome and «cytokine storm» that can lead to death ^8–10^. The penetration of SARS-CoV-2 occurs as a result of the binding of the spike glycoprotein (S-glycoprotein) to angiotensin converting enzyme (ACE2) expressed on the surface of the host cells ^11^, the lung tissues being the main target. Intervention at the stage of adsorption/binding or replication of the virus using therapeutic agents can effectively block viral infection ^12,13^. The most common symptom of COVID-19 patients is respiratory distress, and most patients admitted to intensive care were unable to breathe on their own. In addition, some COVID-19 patients have also experienced neurological signs such as headache, nausea and vomiting. Increasing data show that coronaviruses are not always limited to the respiratory tract and that they can also affect the central nervous system, resulting in neurological complications ^14^. The severe clinical course of the COVID-19 infection indicates the urgent need for research on new antiviral compounds against SARS-CoV-2 ^15,16^.

Animal venoms, containing a wide range of biologically active compounds of different chemical structures, are a rich source of antimicrobial substances ^17–21^. Consequently, animal toxins have significant pharmacological value and great potential for drug discovery ^22–24^. Many snake venom components are being investigated in preclinical or clinical trials for a variety of therapeutic applications ^25–28^. Among the snake venom enzymes, phospholipases A2 (PLA2s) are one of the main components ^29–31^. Typically, PLA2s are small proteins with a molecular weight of ~ 13-15 kDa and belong to the secreted type of enzymes and catalyze hydrolysis at the sn2 position of membrane glycerophospholipids to lysophospholipids and free fatty acids ^32,33^. PLA2s form one of the largest family of snake venom toxins. The whole PLA2 superfamily is currently subdivided into six types, namely secreted PLA2s (sPLA2s), Ca^2+^-dependent cytosolic cPLA2, Ca^2+^-independent cytosolic iPLA2, platelet-activating factor acetylhydrolase PAF-AH, lysosomal LPLA2 and adipose-specific AdPLA2 ^34^. More than one third of the members of the PLA2 superfamily belong to sPLA2, which is further divided into 10 groups and 18 subgroups. The PLA2s from snake venoms are in groups I and II ^35^. PLA2s from Elapidae and Hydrophidae venoms (115-120 amino acid residues and seven disulfide bonds) form the group IA, while those from Crotalidae and Viperidae venoms (120-125 amino acids and seven disulfides) form group IIA. Several PLA2s of group IIA exist as non-covalent dimers formed by enzymatically active subunit and inactive subunit in which Asp residue in the active site is replaced by Gln residue. The examples of such PLA2 dimers are HDP-I and HDP-II from *Vipera nikolskii* venom ^36^. In organism, snake venom PLA2s affect different vitally important system, including nervous system ^37^. It should be noted that endogenous secretory PLA2s play an important role in functioning of the central nervous system in mammals ^38^. PLA2s from snake venoms and from other sources have shown antiviral activity against Dengue and Yellow fever viruses ^39^, HIV ^40^, Adenovirus ^41^, Rous virus ^42^ and Newcastle virus ^43^, demonstrating a great potential of PLA2s as antivirals. It can be expected that PLA2s will have antiviral activity against SARS-CoV-2 and this property can be used in the development of drugs against COVID-19.

Taking this into account, we tested several snake venom PLA2s for their antiviral activity against SARS-CoV-2. We have shown that all tested PLA2s are capable to inhibit SARS-CoV-2 infection *in vitro*, however, only dimeric PLA2s have high antiviral activity. Additional studies on the PLA2s against SARS-CoV-2 showed that the virucidal and antiviral activities of dimeric PLA2 depended on its catalytic activity. The ability of PLA2 to inactivate the virus suggests a new protective mechanism for the host cells that could provide an additional barrier to the spread of infection.

## Results

### Preparation of PLA2s

In total we studied eight samples of snake venom PLA2s and their subunits. Two PLA2s were from krait *Bungarus fasciatus* venom: phospholipase A2 II (GenBank AAK62361.1) and basic phospholipase A2 1 (UniProtKB Q90WA7), named BF-PLA2-II and BF-PLA2-1, respectively. Both these PLA2s belong to group IA. All other PLA2s tested in this work belong to group IIA: one PLA2, Vur-PL2 (UniProtKB F8QN53) was from viper *V. ursinii renardi*. Two more PLA2s, HDP-1 and HDP-2, were from viper *V. nikolskii*. HDP-1 and HDP-2 are non-covalent dimers composed of enzymatically active subunits HDP-1P (UniProtKB Q1RP79) and HDP-2P (UniProtKB Q1RP78), respectively, as well as enzymatically inactive HDP-1I (UniProtKB A4VBF0) which is common for both dimers. By reversed phase chromatography, HDP-2 was separated into HDP-2P and HDP-1I, which were used for further studies. To inactivate HDP-2P, it was treated with 4-bromophenacyl bromide which selectively alkylated His residue in the active site. The modified analogue was purified by reversed phase HPLC and analyzed by mass spectrometry, which indicated the increase in mass by 198 Da (yield of the target derivative was 45%). This increase indicates the inclusion of one 4-bromophenacyl residue into HDP-2P molecule. The study of enzymatic activity showed that phospholipolytic activity of modified HDP-2P decreased by about 2200 times. The inactivated protein called “HDP-2P inact” was used for further studies.

### Antiviral activity of snake venom PLA2s

To study the antiviral activity of PLA2s against SARS-CoV-2, five snake venom PLA2 were tested, two of them, HDP-1 and HDP-2, being dimeric. The antiviral activity of two subunits (HDP-1I and HDP-2P) isolated from HDP-2 was tested as well. In this assay, the cytopathic effect (CPE) of live SARS-CoV-2 on Vero E6 cells was studied. Replication of viruses in Vero E6 cell line produces obvious cytopathic effect in vitro, that is why this cell line is widely used in virology. SARS-CoV-2 infection resulted in morphological changes in cells which included cell rounding, detachment and cytolysis (Fig. 1). We found that all five tested PLA2s showed antiviral activity in an *in vitro* experimental infection model using live SARS-CoV-2 and Vero E6 cells (Fig. 1 and Fig. 2): morphological changes induced by the virus were prevented by PLA2s (Fig. 1). Monomeric group IA BF-PLA2-II (Fig. 1 and Fig. 2) and BF-PLA2-1 (Fig.2) showed low activity, which at the maximum concentration of 100 μg/ml) inhibited SARS-CoV-2 infection by an average of about 50% (Fig. 2). Monomeric group IIA Vur-PL2 had an intermediate antiviral activity: it inhibited infection by 50% at ~1 μg/ml (Fig. 1 and Fig. 2). However, the two investigated dimeric PLA2s HDP-1 and HDP-2 showed a strong antiviral activity, inhibiting CPE even at a concentration of 0.1 μg/ml (Fig. 1 and Fig. 2).

**Figure 1.**
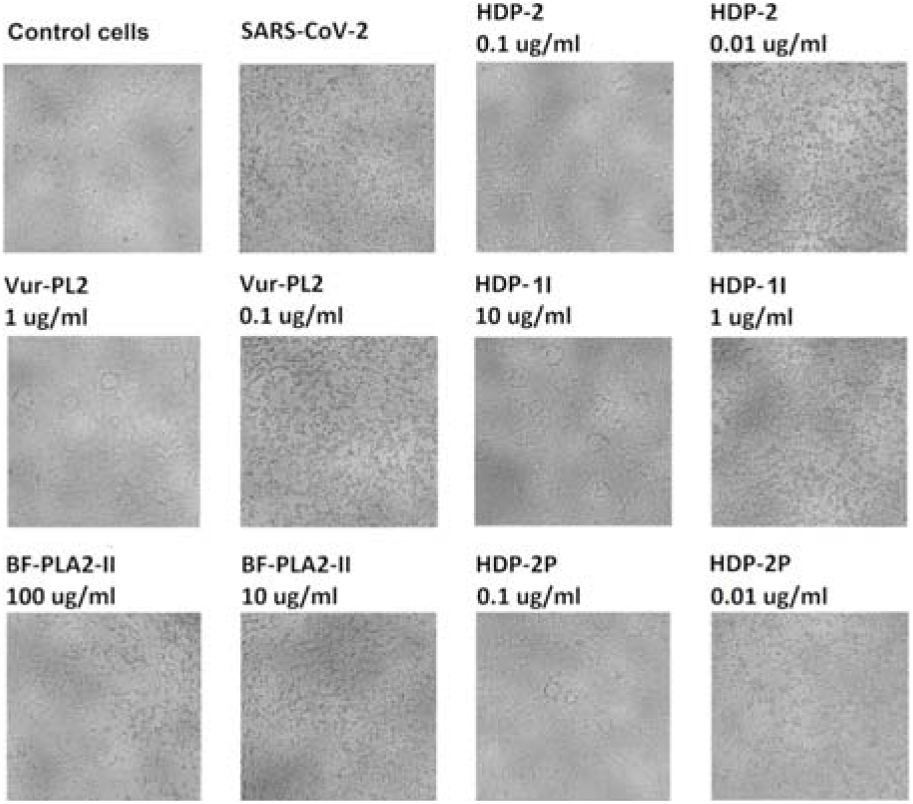
Inhibition of the SARS-CoV-2 cytopathic effect by the snake venom PLA2s. Vero E6 cells were infected with SARS-CoV-2 virus at 100 TCID_50_ (50% tissue culture infectious dose) in the presence of PLA2s at the indicated concentrations. The images were taken 72 h after infection.

**Figure 2.**
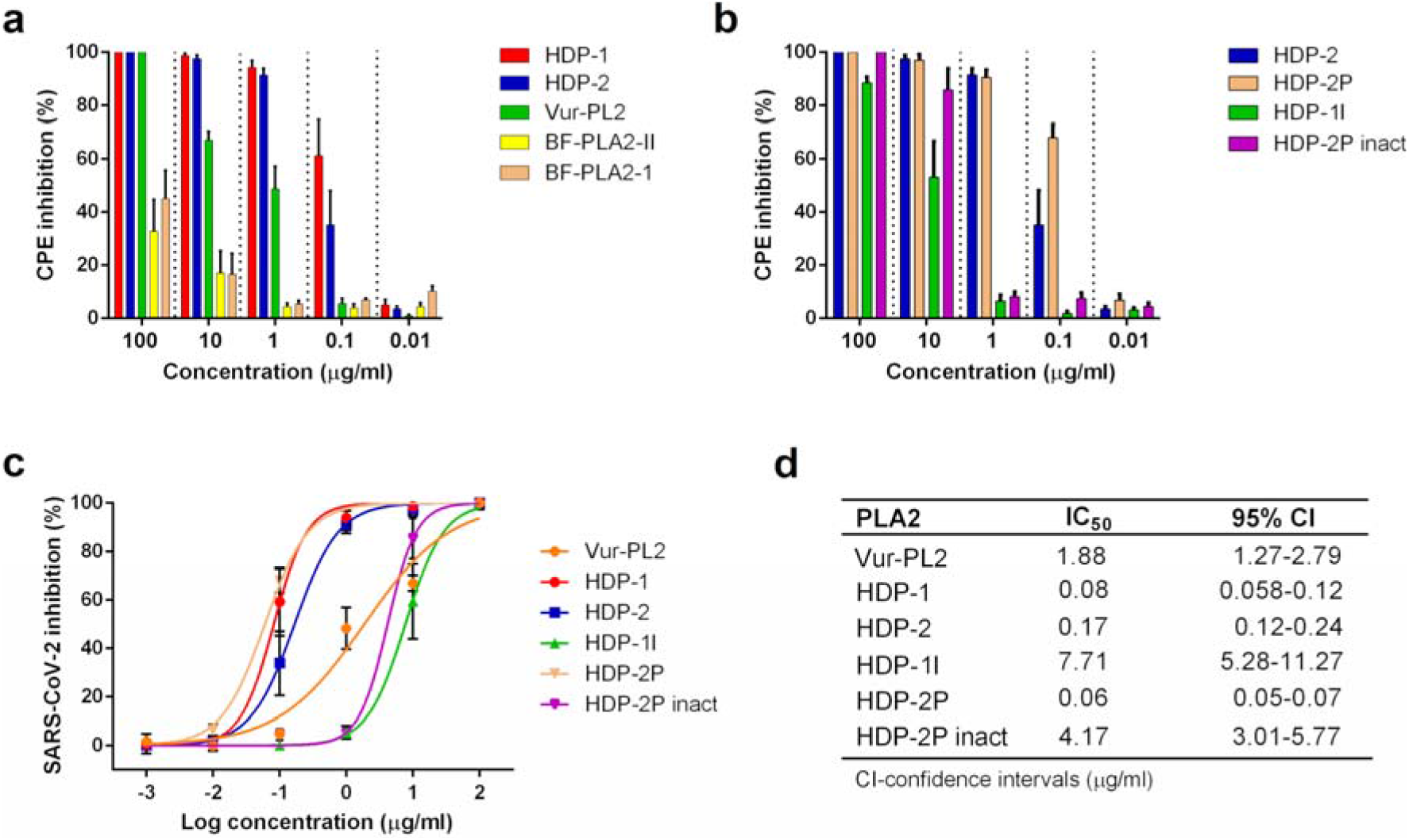
Evaluation of the inhibitory activity of the snake PLA2s against the cytopathic effect of SARS-CoV-2 on the Vero E6 cells. **a** and **b** Vero E6 cells were infected with SARS-CoV-2 at an 100 TCID_50_ in the present of different concentrations of the indicated PLA2s for 72 h. CPE inhibition was then measured by colorimetric assay with 3-(4,5-dimethylthiazol-2-yl)-2,5-diphenyltetrazolium bromide (MTT). **c** Dose - response curves for determination of half-maximum inhibitory concentration (IC_50_) values. Nonlinear regression was used to fit experimental points. **d** IC_50_ values for indicated PLA2s. The results are expressed as mean ± SD of three independent experiments with triplicate measurements.

Next, we studied the antiviral activity of two isolated subunits of HDP-2. The results revealed striking differences between their actions: the catalytically active subunit HDP-2P showed 2-fold higher antiviral activity than the parent HDP-2, while enzymatically inactive subunit HDP-1I showed two orders of magnitude lower antiviral activity than HDP-2 (Fig. 2). This suggests that antiviral activity may be related to phospholipolytic activity. To determine the contribution of the phospholipase activity of the HDP-2P subunit to the inhibition of SARS-CoV-2, the enzymatic activity of this subunit was inhibited by chemical modification of the active site (HDP-2P inact). It was found that after such modification the antiviral activity of HDP-2P decreased 70-fold (Fig. 1 and Fig. 2). Thus, PLA2s from different snakes showed various antiviral activity against SARS-CoV-2. The abolishment of enzymatic activity resulted in a strong decrease of antiviral activity.

### Cytotoxic activity of snake venom PLA2s

To evaluate the direct effects of PLA2s on the Vero E6 cells, the cytotoxicity of PLA2 was studied: it was found that they slightly slow down cell proliferation, without morphological changes in the cell monolayer (Fig. 3). The highest, albeit moderate, cytotoxicity was manifested by Vur-PL2 and HDP-2P which at 100 μg/ml reduced cell proliferation on average by 38% and 51%, respectively. Only HDP-2P at the maximal concentration has pronounced cytotoxicity with a change in cell morphology. However, HDP-2P inhibited SARS-CoV-2 CPE at concentration of 0.1 μg/ml, that is at three order of magnitude lower concentration.

**Figure 3.**
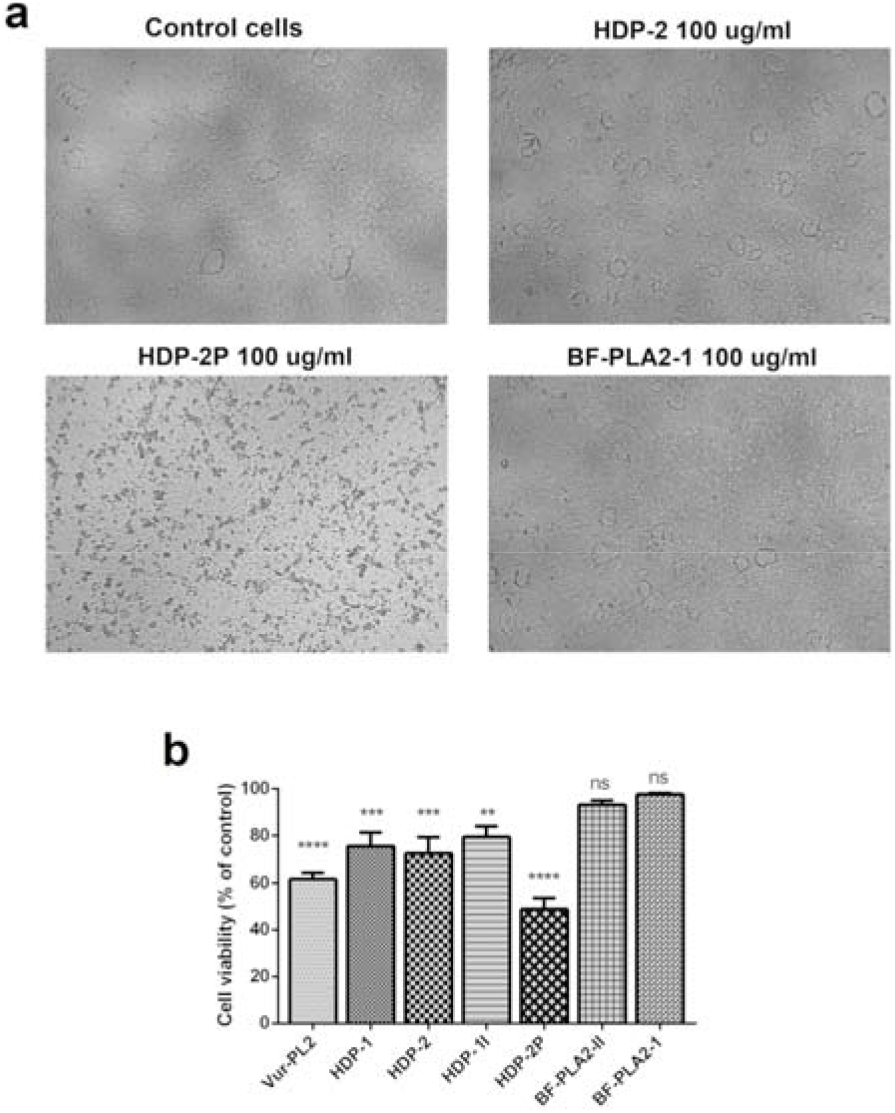
Cytotoxicity of snake PLA2s to Vero E6 cells. **a** The images showing the morphology of Vero E6 cells in the presence of various PLA2s at 100 μg/ml. **b** Cytotoxicity data for all tested PLA2 at concentration 100 μg/ml. Data obtained from three independent experiments with two measurements. The data are presented as the mean ± SD. One-way ANOVA with Turkey post hoc test: ** *p*◻0.01; *** *p*◻0.001; **** *p*◻0.0001; ^*ns*^ *p*>0.05.

### Dimeric PLA2 HDP-2 and its subunits HDP-2P and HDP-1I possesses potent virucidal activity

To elucidate whether the PLA2s exert their antiviral activity through phospholipolytic effects on the cell membrane or on the virus, we analyzed the virucidal activity of the HDP-2 and HDP-2P which manifested the highest antiviral activity as well as of HDP-1I and HDP-2P with the enzymatic activity blocked by chemical modification (HDP-2P inact). To detect a direct inhibitory effect on the viral particles, the SARS-CoV-2 virus stock was treated with various concentrations (0.1–10 μg/ml) of these proteins or buffer. One hour after treatment, the mixtures were diluted 2000-fold and the infectious activity of the virus was determined using serial dilutions method. The results were evaluated after 72 h post infection based on visual scoring of CPE on the Vero E6 cells. Depending on the concentration of HDP-2 and of its subunits, a significant suppression of the infectious viral titer was achieved (Fig. 4). Complete suppression of the infectivity of SARS-CoV-2 was observed when the viral stock was treated with the catalytically active subunit HDP-2P at all tested concentrations. However, as in the experiment on antiviral activity, inhibition of the enzymatic activity of HDP-2P led to a loss in the ability to completely inactivate SARS-CoV-2, which was manifested by an increase in viral titer at lower concentration of HDP-1I or modified HDP-2P.

**Figure 4.**
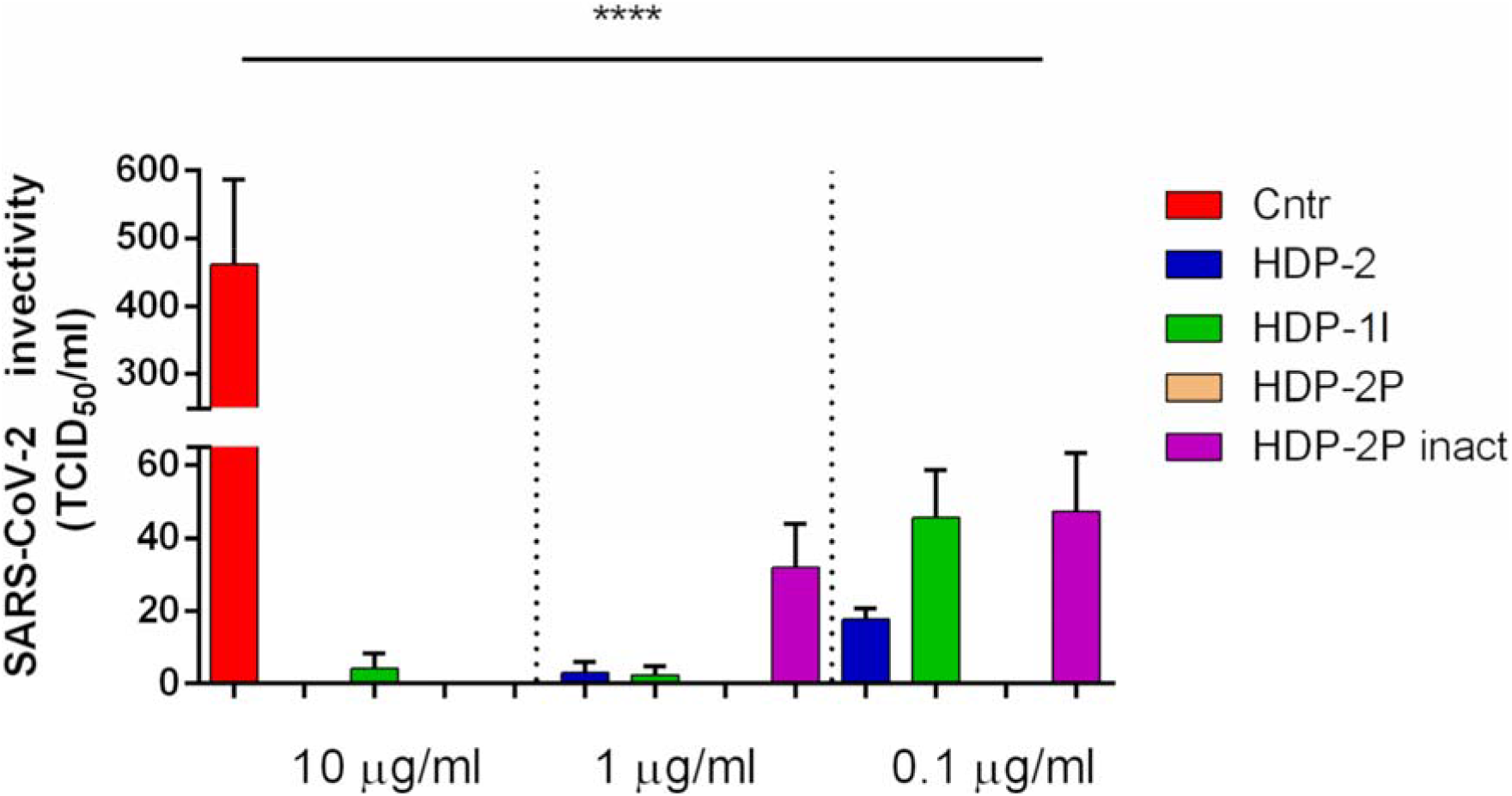
Virucidal activity of HDP-2 and of its subunits. SARS-CoV-2 (1×10^6^ TCID_50_) was incubated with serial decimal dilutions of proteins for 1 h at 37°C. The treated viruses were used to infect the Vero E6 cells. The virus infectivity was calculated by the reduction of the virus titer after treatment with PLA2 in comparison with the untreated virus. The data are presented as mean±SD of three independent experiments and analyzed by one-way ANOVA with Turkey post hoc test. At all protein concentration used *p* < 0.0001 when compared to control. TCID_50_ - 50% tissue culture infectious dose. Cntr - control.

### Inhibition of cell fusion mediated by SARS-CoV-2 glycoprotein S

To study cell fusion mediated by S-glycoprotein interaction with the ACE2 receptor, we used 293T cells expressing green fluorescent protein (GFP) and SARS-CoV-2 S-glycoprotein (293T-GFP-Spike) as well as Vero E6 cells expressing ACE2 which is a receptor for glycoprotein S. The effect of the five snake venom PLA2s on the SARS-CoV-2 S-glycoprotein mediated cell-cell fusion was investigated. After co-cultivation of effector 293T-GFP-Spike cells and target Vero E6 cells at 37°C for 2 h in the presence of various concentrations of PLA2s, the number of fused cells, having the size at least 2 times larger than normal cells and multiple nuclei, were counted using a fluorescence microscope. It was found that at concentrations from 1 to 100 μg/ml the dimeric HDP-1 and HDP-2 inhibited S-glycoprotein mediated cell-cell fusion. They showed approximately the same concentration-dependent activity inhibiting cell-cell fusion by 70% at 100 μg/ml (Fig. 5). At concentration of 1 μg/ml, this value decreased to about 48%, still manifesting statistically significant difference from control. Vur-PL2 and BF-PLA2-II at a concentration of 100 μg/ml showed insignificant inhibition, while HDP-2P at 100 μg/ml completely blocked the cell-cell fusion.

**Figure 5.**
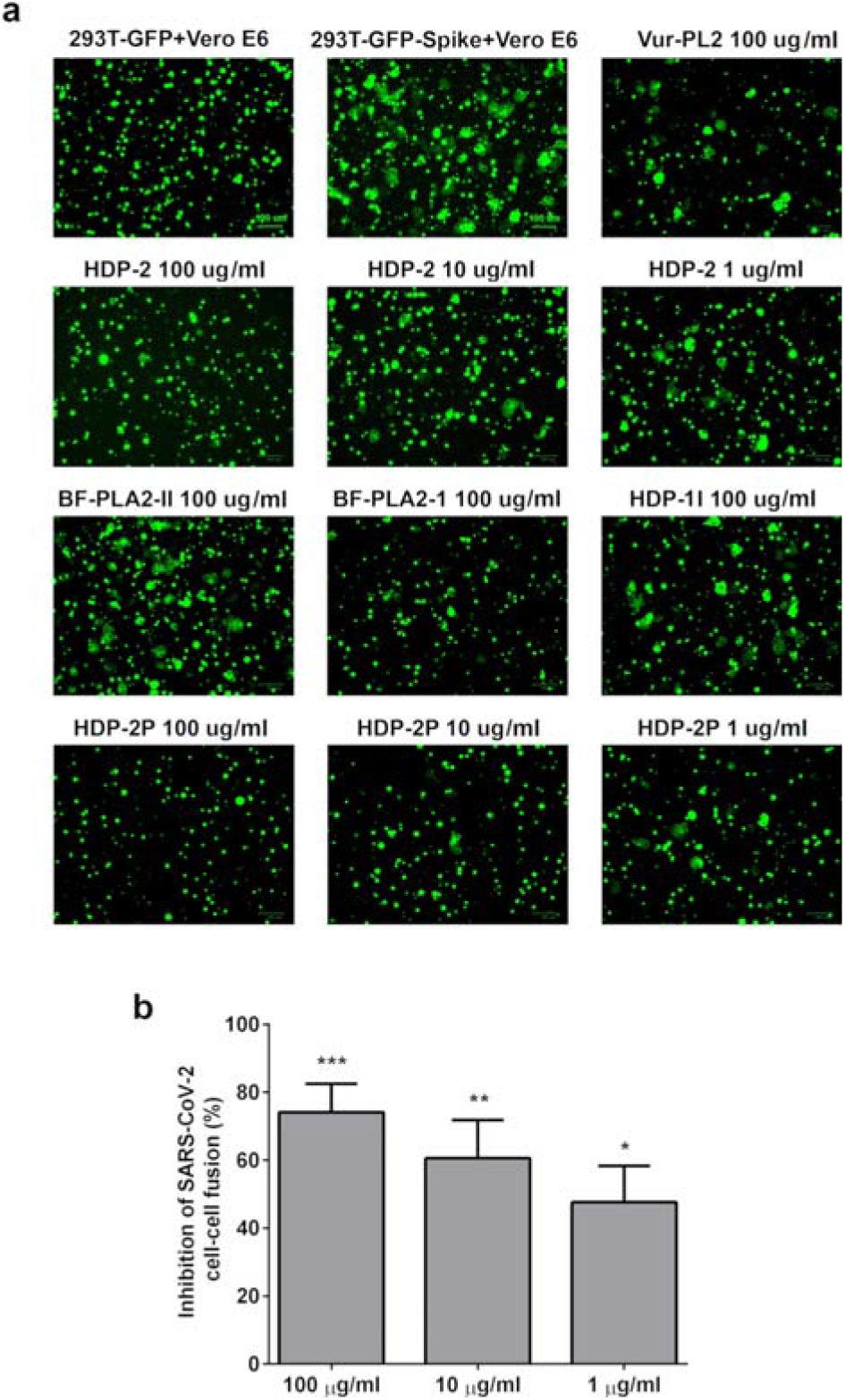
Inhibition of SARS-CoV-2 S-glycoprotein mediated cell–cell fusion by different PLA2s. **a** Images of SARS-CoV-2 S-glycoprotein mediated cell–cell fusion on the Vero E6 cells after 2◻h at different concentrations of PLA2s. **b** The percentage of inhibition of the cell-cell fusion by HDP-2. Cell-cell fusion was calculated relative to the number of fused cells in wells untreated with HDP-2. Experiments were repeated twice, and the data are expressed as means◻±◻SD. One-way ANOVA with Turkey post hoc test: * *p*◻0.05; ** *p*◻0.01; *** *p*◻0.001.

## Discussion

Current efforts to combat the COVID-19 pandemic are primarily focused on hygiene, quarantine of infected people, social distancing and vaccine development ^44,45^. Despite accelerated efforts to develop a vaccine and to find an effective medicine by screening different compounds against COVID-19, patients are in urgent need of therapeutic interventions.

It has been shown that snake venoms have antibacterial ^46^, antiparasitic ^47^, antifungal ^48^ and antiviral ^22^ activities. As discussed in Introduction, snake venoms also represent a promising source of compounds against various viruses. Possessing very diverse sequence variations, sPLA2s show functional variation and are involved in a wide range of biological functions and disease initiation through lipid metabolism and signaling ^49^.

In the present study, we have demonstrated that snake sPLA2s have antiviral activity against SARS-CoV-2. The highest activity was exhibited by dimeric PLA2s of the group IIA. Strong virucidal activity against SARS-CoV-2 was observed when viral particles were treated with HDP-2 or its subunits, especially with enzymatically active subunit HDP-2P. However, the virucidal and antiviral activity of the HDP-2P was markedly suppressed when the His residue in the active center was modified by 4-bromophenacyl bromide, a specific inhibitor of PLA2s. It is interesting to note that a previous study of PLA2 CM-II isoform isolated from cobra *Naja mossambica* venom and belonging to the group IA, showed an almost complete lack of virucidal activity against MERS-CoV ^50^. It should be noted that there are some genetic and structural differences between SARS-CoV-2 and MERS-CoV ^51–56^. For example, the similarity of MERS-CoV glycoprotein S (PDB ID: 5X59) and SARS-CoV-2 S-glycoprotein is about 30%. Given these differences, it can be suggested that PLA2 from different groups can affect the viruses in different ways. These distinctions may be explained by the assumption that some PLA2s selectively degrade the lipid bilayers of the viral envelope through hydrolysis of phospholipids, while others are not able to hydrolyze the viral envelop. The results similar to those obtained in this work for virucidal activity were described in previous studies, where it was shown that dimeric PLA2 crotoxin isolated from *Crotalus durissus terrificus* and its enzymatically active subunit PLA2-CB inactivated Dengue Virus Type 2, Rocio virus, Oropouche virus and Mayaro virus by cleaving the envelope glycerophospholipids that originate from the membranes of host cells ^39,57^. On the contrary, crotoxin and PLA2-CB were inactive against the non-enveloped Coxsackie B5 and encephalomyocarditis virus or viruses budding through the plasma membrane ^57^. It should be noted that the composition of phospholipids in the endoplasmic reticulum membrane differs from the composition of phospholipids in the plasma membrane, which can explain the difference in the composition of phospholipids in the envelopes of different viruses ^58–62^. Depending on the origin and, therefore, the composition of the envelope, PLA2s can differently affect the viruses and exert the different virucidal activity. In addition, several studies support the idea that PLA2s belonging to group V and X inhibit adenoviral infection by hydrolysis of the plasma membrane of host cells ^41,63,64^. The product of PLA2-mediated hydrolysis of phosphatidylcholine, which is part of the membranes, is lysophosphatidylcholine ^65,66^ which has been shown to inhibit membrane fusion caused by influenza, SIV, Sendai and rabies viruses, blocking viral entry into host cells ^67–70^. All PLA2s investigated in this work showed only weak cytotoxicity against Vero E6 cells, which was manifested by a decrease in cell proliferation. It should be noted that we and other researchers have previously observed antiproliferative activity of PLA2s against cancer cells ^71,72^. However, it is important to emphasize that PLA2s have a cytotoxic and antiproliferative effect mainly on cancer cells without affecting normal cells ^28,71,73^. Thus, considering that PLA2s studied in this work inhibit SARS-CoV-2 at concentrations several order of magnitude lower than those affecting normal cells, they can be regarded as potential basis for design of antiviral drugs or as valuable tools for the study of virus interaction with host cells.

The envelopes of coronaviruses contain spike glycoproteins S which form a “crown” and mediate the viral entry into the cells. Glycoprotein S is composed of two units (domains): S1, which binds to the ACE2 receptor on the surface of host cells, and S2 which mediates fusion of the viral envelope with the cell membrane. In a virus not associated with a cell, the glycoprotein S present on the surface of the SARS-CoV-2 is inactive. However, after binding to the cellular receptor ACE2, target cell proteases activate the glycoprotein S, cleaving site between S1 and S2 and leaving the S2 subunit free to mediate viral fusion and penetration ^74,75^. SARS-CoV-2 can enter the target cell in two ways: one is the pathway of endocytosis, and the other is direct fusion at the cell surface ^76,77^. We found that dimeric PLA2 HDP-1 and HDP-2 are able to block SARS-CoV-2 glycoprotein S mediated cell-cell fusion. Thus, we can hypothesize that dimeric PLA2s are capable of blocking viral entry into cells like peptide fusion inhibitors ^78,79^, manifesting an additional mechanism of antiviral activity. It should be noted that a number of peptides from scorpion venom have been shown to be active against such retroviruses as HIV/SIV through their ability to bind to the HIV glycoprotein gp120 due to molecular mimicry of the CD4^+^ receptor. As a result, they block the gp120-CD4 interactions, which are important for the initiation of conformational changes in the viral envelope that trigger virus entry into the host cells ^80^. To check the homology between glycoprotein S and dimeric PLA2s, we have performed the alignment of the glycoprotein S amino acid sequence with those of HDP-2 and HDP-1. Some similarity between the sequences was observed (Fig. 6). Interestingly, the amino acid sequences of PLA2s have similarity to the glycoprotein S fragment 413-462, which according to the X-ray structure is involved in interaction with ACE2 ^81^. Amino acid residues Lys417, Tyr449 and Tyr453 forming contacts with ACE2 are conserved in PLA2 sequences. Considering this homology, we suggest that dimeric PLA2s may compete with SARS-CoV-2 for binding to ACE2. However, a true mechanism for blocking S-glycoprotein mediated cell-cell fusion by PLA2s remains to be studied.

**Figure 6.**
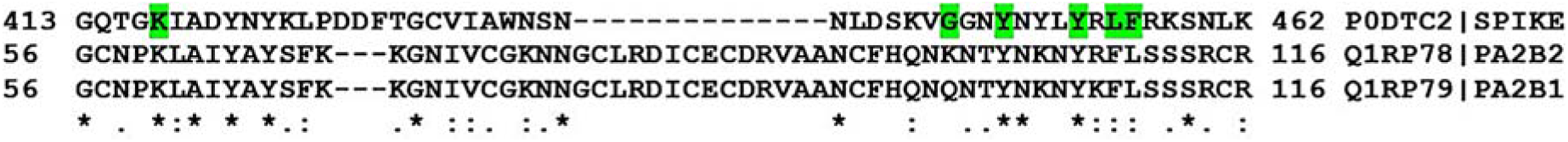
Alignment of the amino acid sequence of SARS-CoV-2 glycoprotein S with sequences of enzymatically active subunits of dimeric PLA2s from *V. nikolskii*. The amino acid residues of glycoprotein S forming contacts with ACE2 are marked in green. P0DTC2 – S-glycoprotein of SARS-CoV-2, Q1RP78 – HDP-2P, Q1RP79 - HDP-1P. The alignment was done using on-line multiple alignment program MAFFT (version 7) at Computational Biology Research Consortium (CBRC, Japan)^89^.

In this study, we have demonstrated for the first time the antiviral activity of snake PLA2s against SARS-CoV-2. Dimeric HDP-2 and its catalytic subunit showed high virucidal activity, most likely due to the cleavage of glycerophospholipids on the virus envelope, which can lead to the destruction of the lipid bilayer and destabilization of surface glycoproteins in the virus. Summarizing previous and our findings on the ability of snake PLA2s to inactivate various viruses, these results highlight the potential of PLA2s as a natural product for the development of broad-spectrum antiviral drugs.

## Materials and Methods

### Cells, virus and plasmid

Vero E6 (ATCC CRL-1586) and 293T (ATCC CRL-3216) cells were grown in complete DMEM medium (Gibco, USA), supplemented with 10% fetal bovine serum (FBS; HyClone, USA), 1× L-Glutamine (PanEco, Russia) and 1× penicillin-streptomycin (PanEco, Russia). All cell lines were tested negative for mycoplasma contamination. SARS-CoV-2 strain hCoV-19/Russia/Moscow_PMVL-4 (GISAID ID: EPI_ISL_470898) was isolated from naso/oropharyngeal swab of COVID-19 patient using the Vero E6 cell line. The stocks of SARS-CoV-2 used in the experiments had undergone six passages. Viral titers were determined as TCID_50_ in confluent Vero E6 cells in 96-well microtiter plates. Virus-related work was performed at biosafety level-3 (BSL-3) facility. pUCHR-IRES-GFP plasmid was kindly provided by D.V. Mazurov (Institute of Gene Biology, Moscow, Russia). pVAX-1-S-glucoprotein plasmid encoding a full-length SARS-CoV-2 Spike glycoprotein (Wuhan) was obtained through Evrogen (Russia).

### Phospholipases A2

Phospholipase A2 II (BF-PLA2-II, GenBank AAK62361.1) and phospholipase A2 1 (BF-PLA2-1, UniProtKB Q90WA7) were purified from krait *Bungarus fasciatus* venom as described ^71^. Phospholipase A2 Vur-PL2 (UniProtKB F8QN53) was purified from viper *V. ursinii renardi* venom as described in ^82^. Dimeric phospholipases HDP-1 and HDP-2 were isolated from vipera V. nikolskii venom and separated into subunits HDP-1P (UniProtKB Q1RP79), HDP-2P (UniProtKB Q1RP78) and HDP-1I (UniProtKB A4VBF0) as described in ^36^.

### Modification of HDP-2P

4-Bromophenacyl bromide (Lancaster, England) as 10 mM stock in acetone was added to final concentration of 200 μM to the 20 μM HDP-2P solution in 50 mM Tris HCl buffer, pH 7.5, containing 10 mM Na_2_SO_4_. The mixture was incubated for 6 h at a room temperature and separated using a Jupiter C18 HPLC column (Phenomenex) and acetonitrile gradient from 20 to 50% in 30 min in the presence of 0.1% trifluoroacetic acid. The phospholipolytic activity was measured according to ^83^ using a synthetic fluorescent substrate 1-palmitoyl-2-(10-pyrenyldecanoyl)-sn-glycero-3-phosphocholine (Molecular Probes, The Netherlands) and a Hitachi F-4000 spectrofluorimeter.

### Cell viability assay

Cell viability in the presence of the tested PLA2 was assessed using the MTT (Methylthiazolyldiphenyl-tetrazolium bromide) method as previously described ^84^. Briefly, Vero E6 cells (2 × 10^4^ cells per well) were incubated with different concentrations of the individual PLA2s in 96-well plates for 48 h, followed by the addition of the MTT solution (Sigma, USA) according to the manufacturer’s instructions. Results were expressed as percentage of cell viability in comparison to the untreated cell controls. Three experiments were performed with two technical replicates.

### Cytopathic effect (CPE) inhibition assay against SARS-CoV-2

Vero E6 cells were plated in 96-well plates at a density of 2 × 10^4^ cells per well. After 18 h of incubation, 100 TCID_50_ SARS-CoV-2 were added to the cell monolayer in the absence or presence of various concentrations of PLA2s (10-fold dilutions in a concentration range of 0.001 μg/ml to 100 μg/ml). After 72 hours of incubation, differences in cell viability caused by virus-induced CPE were analyzed using MTT method as previously described ^85,86^. For this, an MTT stock solution (5 mg/ml in PBS) was added to each well at a final concentration of 0.5 mg/ml. After 2 h incubation, medium was aspirated from wells and 150 μl of DMSO was added. Absorbance was measured at 590 nm using SPECTROstar Nano microplate reader (BMG LABTECH). Three independent experiments with triplicate measurements were performed. Data were analyzed using GraphPad Prism 6 (GraphPad Software Inc., La Jolla, CA, USA) and the IC_50_ (concentration of the compound that inhibited 50% of the infection) were calculated via nonlinear regression analysis.

### Virucidal activity test

The SARS-CoV-2 virus stock (1×10^6^ TCID_50_) was incubated with serial decimal dilutions of PLA2 (0.1-10 μg/ml) for 1 h at 37°C. Then, the virus samples treated by PLA2 were diluted below IC_50_ and titrated on Vero E6 cells by limiting dilution assay (5-fold dilutions in six replicates) in 96-well plates. A viral stock treated with PBS was used as a control. The plates were incubated at 37°С (5% CO_2_) for 72 hours. The CPE was scored visually under a microscope and the virus titers were calculated by the Reed and Muench method ^87^.

### Cell-Cell fusion assay

The inhibitory activity of PLA2 on a CoV-2 S-mediated cell–cell fusion was assessed as previously described ^79,88^. 293T effector cells were transfected with plasmid pUCHR-IRES-GFP encoding the GFP and plasmid pVAX-1-S-glucoprotein encoding CoV-2 S protein (293T-GFP-Spike). Vero E6 cells (3×10^4^ cells per well), expressing ACE2 receptors on the membrane surface, were used as target cells, being incubated in 96-well plates for 18 h. Afterwards, 10^4^ effector cells (293T-GFP-Spike) per well were added in the presence or absence of PLA2 at various concentrations and incubated at 37◻°C for 2◻h. The percentage of cell-cell fusion was calculated by counting the fused cells in each well in five random fields using an Olympus (Japan) epifluorescence microscope.

### Statistical analysis

Data was analyzed by one-way ANOVA with a Tukey’s post hoc test for multiple comparisons. *P*◻<◻0.05 is considered statistically significant. All analyses were performed with GraphPad Prism version 6.

## Acknowledgements

This work was supported by the Russian Foundation for Basic Research (RFBR) grant No. 20-04-60277.

## Author Contributions

A.E.S. designed and performed the experiments; S.D.G. and M.A.N. performed viral experiments and helped in data collection; A.V.O., V.G.S. and Yu.N.U. prepared compounds and performed enzyme activity assays; V.A.G. helped with data interpretation; V.I.T. and Yu.N.U. supervised the project and editing of the manuscript. All authors analyzed the results and approved the final version of the manuscript.

